# Host immunity alters successional ecology and stability of the microbiome in a C. elegans model

**DOI:** 10.1101/2020.06.26.174706

**Authors:** Megan Taylor, NM Vega

## Abstract

A growing body of data suggests that the microbiome of a species can vary considerably from individual to individual, but the reasons for this variation - and the consequences for the ecology of these communities – remain only partially explained. In mammals, the emerging picture is that the metabolic state and immune system status of the host affects the composition of the microbiome, but quantitative ecological microbiome studies are challenging to perform in higher organisms. Here we show that these phenomena can be quantitatively analyzed in the tractable nematode host *Caenorhabditis elegans*. Mutants in innate immunity, in particular the DAF-2/Insulin Growth Factor (IGF) pathway, are shown to contain a microbiome that differs from that of wild type nematodes. We analyze the underlying basis of these differences from the perspective of community ecology by comparing experimental observations to the predictions of a neutral sampling model and conclude that fundamental differences in microbiome ecology underlie the observed differences in microbiome composition. We test this hypothesis by introducing a minor perturbation to the colonization conditions, allowing us to assess stability of communities in different host strains. Our results show that altering host immunity changes the importance of inter-species interactions within the microbiome, resulting in differences in community composition and stability that emerge from these differences in host-microbe ecology.

**Importance:** Here we use a *Caenorhabditis elegans* microbiome model to demonstrate how genetic differences in innate immunity alter microbiome composition, diversity, and stability by changing the ecological processes that shape these communities. These results provide insight into the role of host genetics in controlling the ecology of host-associated microbiota, resulting in differences in community composition, successional trajectories, and response to perturbation.

## Introduction

Host-associated microbiomes are increasingly recognized as ecological systems, where interactions among microbes and between microbes and their host are important for shaping community composition, structure, and function (1, 2). As this understanding has developed, there has come a search for the ecological principles that define these communities (3, 4).

There is particular interest in understanding the sources and consequences of variation in host-associated microbiota. Host-associated microbiomes associated with any given body site can vary considerably between individuals and within individuals over time (5, 6). Some of this variation is attributable to the stochastic processes of colonization and drift experienced by any open ecological system (7–11). However, there is a gathering consensus that host-associated microbial communities are shaped by deterministic processes, including filtering (selection of colonists by the host) and competitive and cooperative interactions among microbes (10, 12, 13).

Some of this variation can be traced to genetic differences between hosts (14–16). However, the effect of host genetic variation on the ecology of microbiome communities remains poorly understood. Much of the existing data focuses on interactions between individual commensal bacteria and their specific hosts, although there is increasing interest in understanding the mechanisms by which a host can control the composition of its microbiome (17–19). If host genetics affect the ecological processes of community assembly, differences between hosts can result in differences in microbiome succession, composition, and stability properties due to differences in the underlying ecology of these communities. Understanding how the host environment shapes the ecological dynamics of microbiome communities will be important for determining how communities in different hosts might respond differently to normal perturbations (e.g. changes in nutrient availability, exposure to innocuous microbes, host circadian rhythms) and pathological events (e.g. pathogen invasion, drug or toxin exposure, host disease or trauma).

Here we present experimental evidence that differences in host genetics can alter the ecological dynamics of microbiome communities, resulting in differences in assembly, succession, and response to perturbation. Using a minimal native microbiome of the nematode *Caenorhabditis elegans*, we colonized N2 wild-type hosts and well-characterized mutant strains under highly controlled conditions to determine the effects of host genetics on the structure and dynamics of intestinal communities.

## Results

In these experiments, germ-free, reproductively sterile adult *C elegans* from N2 (wild type) and selected mutant strains (**Table 1**) were colonized from an artificially constructed metacommunity of eight bacterial strains. These bacterial strains were derived from a wild *C. elegans* microbiome (20) (**Methods**) and were selected to represent the overall diversity of bacterial taxa in these samples and also because each possessed a unique colony morphology when co-cultured on agar plates (**Supp Fig 1**).

**Table 1.** *C. elegans* lineages used in this study. Pharyngeal mutants are shown in blue; innate immune mutants are in yellow; control strains are in grey.

| Worm Strain | Mutation | Description |
| --- | --- | --- |
| N2 |  | Wild type |
| DA573 | eat-14(ad573) X |  |
| DA597 | phm-2(ad597) I |  |
| CF1038 | daf-16(mu86) I | DAF-2/IGF defective |
| CB1370 | daf-2(e1370) III | DAF-2/IGF up-regulated |
| NU3 | dbl-1(nk3) V | TGF-β defective |
| BW1940 | ctls40 [dbl-1(+) + sur-5::GFP] | TGF-β over-expression |
| AU37 | glp-4(bn2) I; sek-1(km4) X | p38 MAPK defective |
| JT366 | vhp-1(sa366) II | p38 MAPK over-expression |
| SS104 | glp-4(bn2) I | AU37 control strain |

Bacterial communities in the N2 intestine showed distinct ecological patterns despite high variability (**Fig. 1, Table S1**). While total colonization ranged across nearly two orders of magnitude, from 1,680 to 67,200 CFU/worm, community composition did not change dramatically across this range (**Fig. S2**). Overall, these communities were characterized by high frequencies of MYb71 (*Ochrobactrum*) (**Fig. 1B**), consistent with prior observations that this genus tends to dominate communities in lab-colonized worms (20). Community composition changes over the course of colonization (**Fig. 1C**), and day 4 of colonization appears to represent mid-succession in these experiments. We therefore opted to measure communities after four days, which we hypothesized would provide useful data on differences in ecological succession across host strains, albeit while failing to maximize compositional differences between lineages.

**Figure 1.**
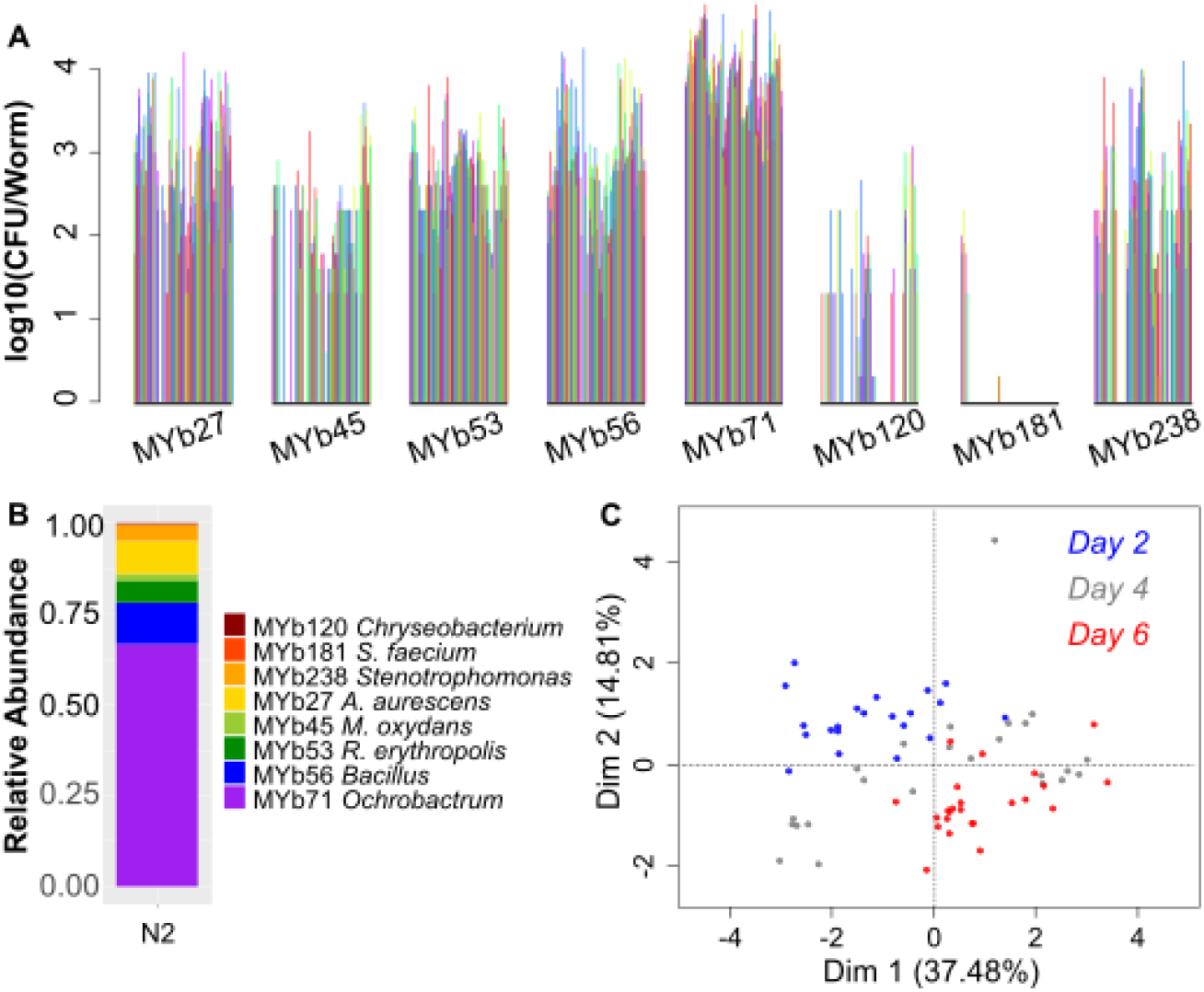
Eight-species microbial communities in the N2 intestine show distinct trends and variation. (A) CFU/worm data for the full data set of individual hosts (n=164) after four days of colonization with this eight-species bacterial consortium; each color represents a single host, and data are grouped by bacterial species to illustrate trends in abundance. (B) Average relative abundance of each bacterial species across all individual N2 worms. (C) Principal component analysis (PCA) of community composition over time in N2 worms across a six-day time series of colonization (24 individual worms/day, destructive sampling of individual hosts).

Next, we explored the effects of host mutations on the composition of the microbiome (**Fig. 2, Table S1-2, Fig. S3**). All worm mutants were colonized under the same conditions used for N2 (above). First, we analyzed grinder-defective mutants (*eat*, also *phm-2*), which have known alterations in their interactions with bacteria (21, 22) but previously unexplored effects on bacterial community composition. Bacterial communities in the mild grinder mutant *eat-14* were very similar to those seen in N2. While communities in the severe grinder mutant *phm-2* were large compared to N2 (median CFU/worm 36,190 vs 13,000), composition was similar to N2 communities of the same size (**Fig. S3**), indicating that the increased permissiveness of the defective grinder did not substantially affect community assembly.

**Figure 2.**
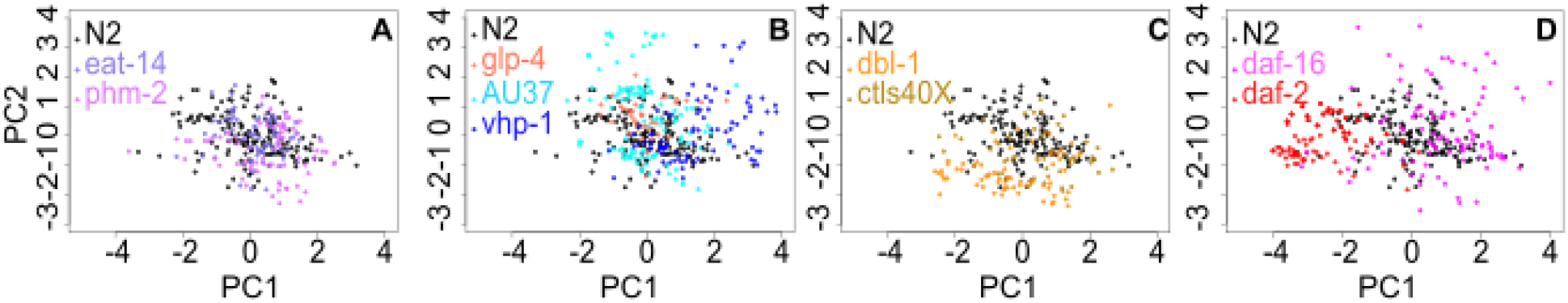
PCA of intestinal community composition in N2 wild-type (n=164, black points) and mutant hosts colonized from a uniform metacommunity of eight bacterial species from the C. elegans native microbiome. (A) Mild (*eat-14*, n=69) and severe (*phm-2*, n=78) grinder mutants, (B) p38 MAPK pathway defective (AU37, n=115) and de-repressed (*vhp-1*, n=84) mutants, with *glp-4* (n=36) control for the AU37 strain; (C) TGF-β defective mutant (*dbl-1*, n=69) and over-expression construct (*ctls40*, n=48), (D) DAF-2/IGF defective (*daf-16*, n=100) and de-repressed (*daf-2*, n=98) mutants. All data represent the results of single worm digests and plating after four days total colonization on the uniformly distributed synthetic eight-species bacterial consortium.

We then explored the effect of the well-conserved innate immune system of *C. elegans* (23, 24) on composition of the microbiome. We found that bacterial communities in innate immunity mutants showed differences from wild type N2 (**Fig. 2B-D, Fig. S3**), but the relationship between community composition and immune function in this host is complex. Neither gain or loss of function in innate immune pathways had a consistent effect on microbiome composition. As previously described (25), a loss of function in the p38 MAPK pathway (AU37) produced microbial communities that closely resembled those in a *glp-4(bn2ts*) control and in N2; however, de-repression of this pathway (*vhp-1*) resulted in a microbiome that differed from N2 (**Fig. 2B**). Conversely, loss of function in the TGF-β pathway (*dbl-1*) produced communities that diverged from wild type as previously observed (25), while over-expression (*ctls40*) did not seem to affect microbiome composition (**Fig. 2E**). Finally, both loss of function (*daf-16*) and derepression (*daf-2*) of the DAF-2/IGF pathway produced marked changes in microbiome composition and variation as compared with N2 (**Fig. 2F**).

Next we sought to establish ecological mechanisms underlying the observed differences in microbiome composition. As previously indicated, we chose to sample communities after four days of colonization to capture mid-successional ecology. N2 wild-type worms showed a negative trend in the relationship between diversity (measured as Shannon H) and total microbiome size (CFU/worm in individual worms) (**Fig S4A**), although the considerable compositional and structural variation in these communities (**Fig. 1**) results in a poor overall fit for the simple linear model (**Table S3**). This is consistent with canonical ideas of competitive succession in ecological communities, where less crowded communities (representing earlier succession) show high diversity, which decreases as the environment fills and competition for resources intensifies. Pharyngeal mutants showed diversity-size relationships similar to N2 (**Fig S4B**), consistent with the compositional similarities observed (**Fig. 2**). Among the innate immune system mutants (**Fig. 3A-C, Table S4**), most of the lineages that showed compositional differences from N2 also showed differences in the diversity-size relationship (26) (*vhp-1* with likelihood ratio test *p*=1.94*10^−4^, **Fig. 3A**; *daf-2 p*=5.55*10^−7^ and *daf-16 p*=1.61*10^−9^, **Fig. 3C**; **Table S3**), suggesting that differences in competition between gut bacteria and the process of ecological succession underlie the observed differences in community composition. (Note that *dbl-1* adults are physically smaller than N2, which likely explains the smaller populations observed in these hosts (27). Also, note that several of these linear fits indicate no significant relationship between Shannon diversity and total microbiome size, and the large β-diversity within host lineages is expected to be a factor here.)

**Figure 3.**
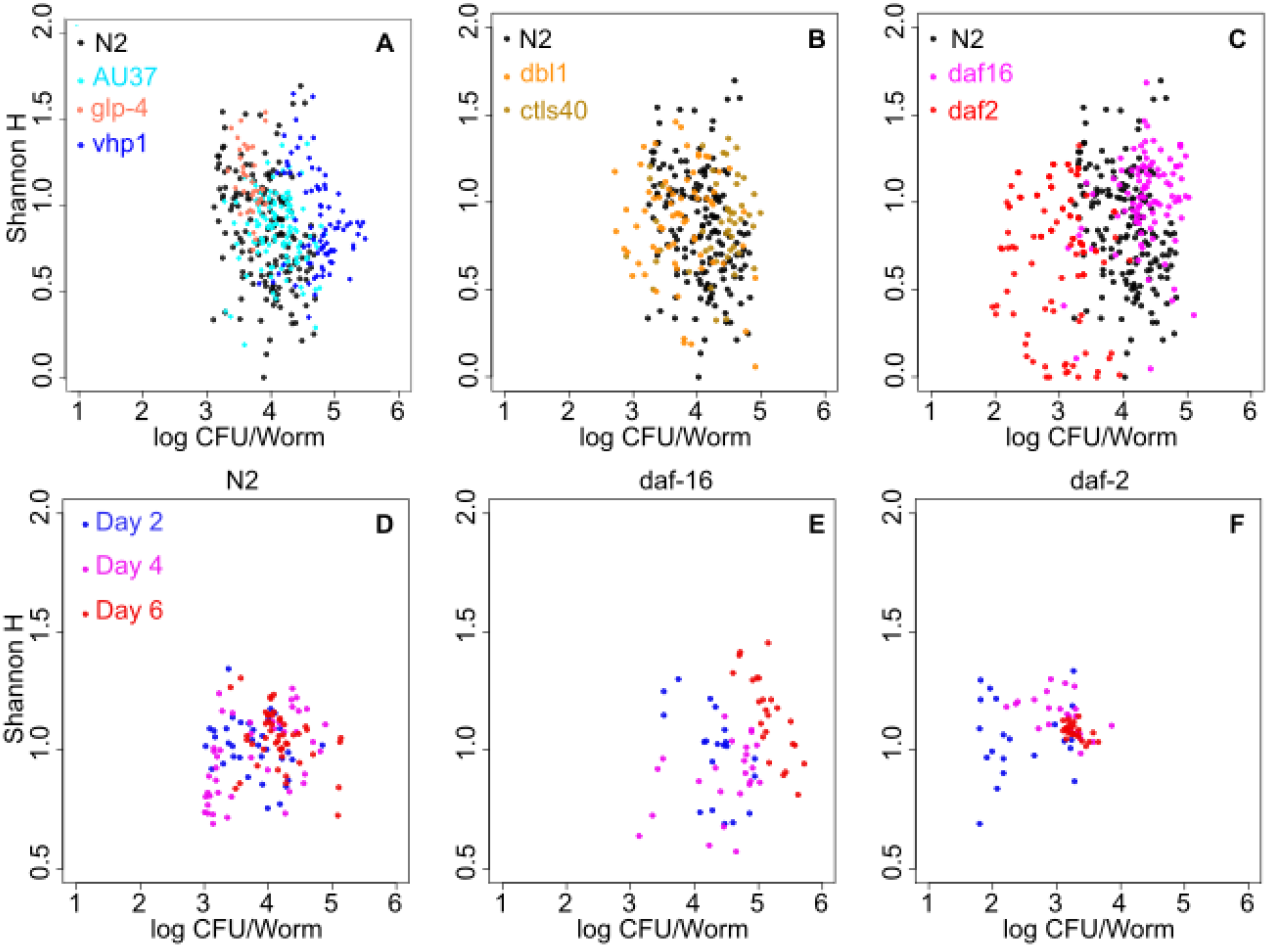
Successional ecology is altered by host genetic background. (A-C) The diversity-population size relationship in intestinal bacterial communities differs across host lineages. Mutants in the (A) p38 MAPK, (B) TGF-B, and (C) DAF-2/IGF pathways show trends that diverge in some cases (see Table S2) from that observed in N2 (black dots). Note that Shannon diversity in these experiments has a maximum at ln(8)=2.08. (D-F) Time series of the diversity-population size relationship indicate that succession is altered in DAF-2/IGF mutant hosts. Intestinal communities were quantified for individual worms (n=24 worms per time point per condition) of (D) N2, (E) *daf-16*, and (E) *daf-2* lineages after two, four, and six days of colonization on the uniform eight-species metacommunity.

If ecological succession differs between *C. elegans* mutant strains, there should be differences in microbiome development over time. Due to the magnitude and bi-directional nature of the divergences from wild-type, we chose to use DAF-2/IGF mutant hosts in these experiments. Here we colonized adult worms from the N2, *daf-16*, and *daf-2* lineages on the uniform eight-species metacommunity and quantified intestinal communities at 48-hour intervals during community development. In N2 hosts, we observe convergence of communities over time as previously described (**Fig. 1C**); by day six of community development, N2-associated intestinal communities had largely converged to a fairly wide but well-defined range (10^3.5-4.5^ CFU/worm, 0.5<H<1.5), and the negative diversity-population size relationship had diminished, suggesting a later stage of succession (**Fig. 3D**, **Fig. S5A-B**). *daf-16* hosts displayed large populations which continued to increase in size and diversity over the observed period (**Fig. 3D, Fig. S5C-D**), while *daf-2* hosts showed progressively less complex microbiomes consisting mainly of three dominant bacteria (**Fig. 3E, Fig S5E-F**). These data (**Fig. 3D-F, Fig. S5**) show that differences in ecological succession underlie the observed differences in community composition across *C. elegans* mutant strains. However, we hypothesized that differences in bacterial interactions underlie differences in microbiome successional ecology, and these experiments do not directly test for this mechanism.

Genetic differences between hosts could alter microbiome ecology and succession by altering the efficiency of host selection during colonization (environmental filtering), by changing the inter-species interactions among bacteria, or both. This should be associated with differences in inter-species correlations within these communities. In this scenario, the *daf-2* mutant would represent a highly selective environment; due to strong interactions with the host, bacterial density never rises to the level where competition between strains is a major influence. The anticipated outcome would be strong compositional convergence enforced by environmental filtering and weak negative correlations among bacterial species. Conversely, the *daf-16* mutant would represent an environment with poor active selection by the host. A lack of host control could produce the large and diverse communities observed, and should result in stronger than expected correlations (negative and/or positive) among bacterial strains due to increased interaction in the intestine.

To test this hypothesis, we first calculated the Spearman correlations among intestinal bacterial strains for each host strain, using the large data set underlying **Figure 2**. Intestinal communities in all host lineages showed a range of correlations, and the distributions of inter-species correlations differed across hosts. The *daf-16* communities showed a broad distribution of correlations relative to N2, with fewer near-zero interactions, while *daf-2* communities showed stronger positive correlations but weaker negative correlations than in N2 (**Fig. 4A-C, Fig. S6**).

**Figure 4.**
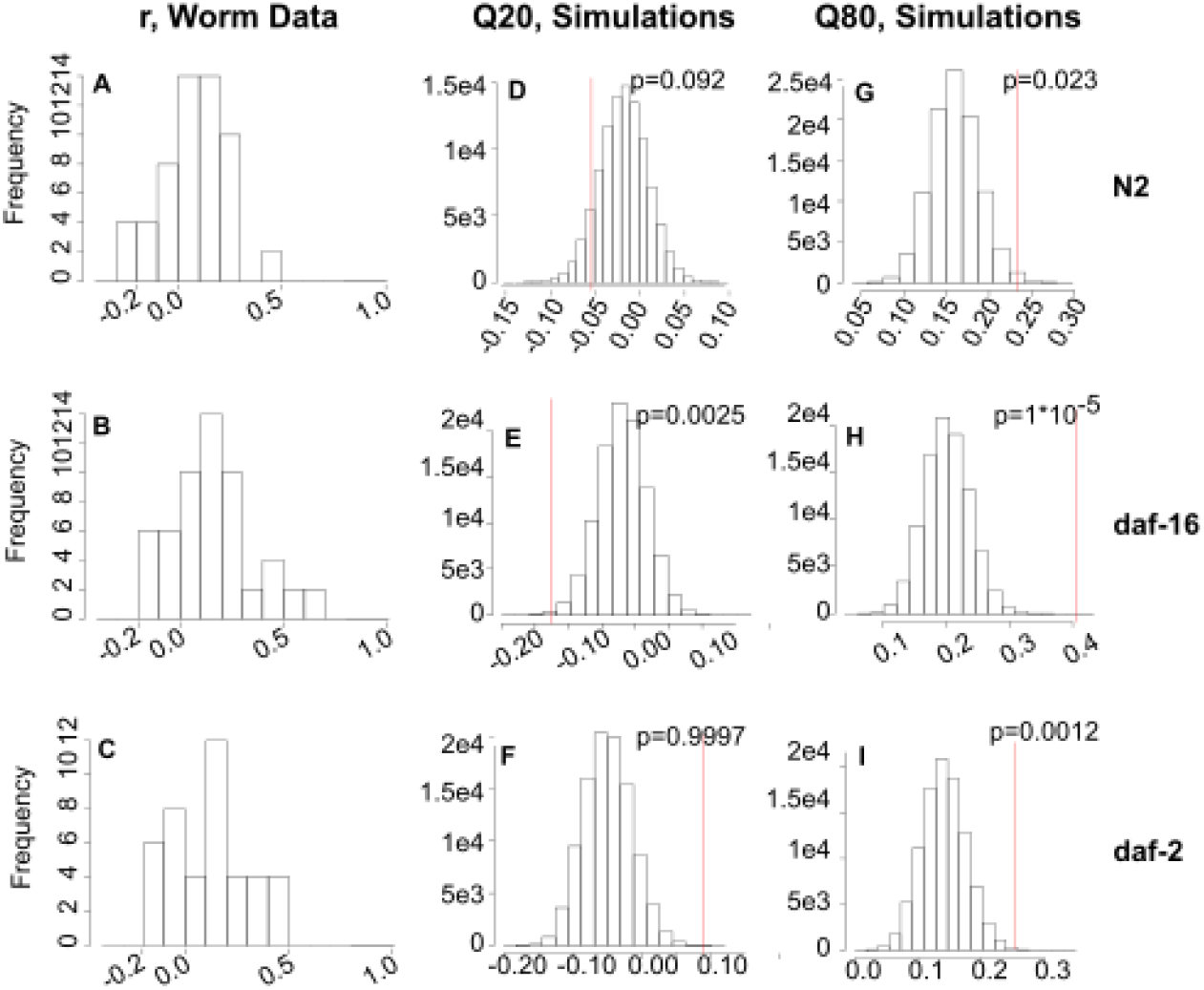
Spearman correlations between bacteria in host-associated intestinal communities differ from the predictions of a neutral sampling model. (A-C) Histograms of Spearman correlation coefficients calculated from data for (A) N2 (n=164), (B) daf-16 (n=100), and (C) daf-2 (n=98) intestinal communities, bin size 0.1. (D-I) Histograms of Spearman coefficient 20^th^ and 80^th^ quantiles from data simulated using host lineagespecific parameterizations of the Dirichlet-multinomial model (n=10,000 simulated data sets per condition). Red lines indicate the corresponding quantile from the empirical data, and p-values indicate the one-tailed percentage of simulated data sets with a lower 20^th^ quantile (D-F) or higher 80^th^ quantile (G-I) than the empirical data.

We next sought to quantify expectations for these distributions that take into account differences in sample size between lineages and in community size between individuals. As a full stochastic model of colonization would require estimation (empirical or otherwise) of a fairly large number of parameters, we opted for a simpler data-driven approach where we fit each data set to a neutral sampling distribution.

The Dirichlet-multinomial distribution is a multivariate sampling distribution that has frequently been used to describe community data of this kind (28–30). In the neutral sampling process described by this model, bacterial species are allowed to have different probabilities of being “drawn” from the pool of potential colonists (in this scenario, where all species are present at equal levels in the metacommunity, this corresponds to different effective migration rates into the host) but species do not differ in their ability to fill space (compete) within the host. By comparing the empirical distributions of correlations to those produced under the neutral sampling assumption, we can determine whether our hypothesis – that the host immune system modifies the effective strength of interactions between bacteria in the gut, possibly by changing the relative importance of interactions with the host – is supported.

We fit this model to our data (**Table S4**) and used these parameterized distributions to generate simulated communities that replicate the structure of the real data, with the same number of hosts and the same number of bacteria in each host (**Fig. S7**). By generating a large number of simulated data sets (n=10,000) for each host lineage, we can compare the correlations observed between bacterial species in real communities to those obtained from similar-quality simulated data where there are no true interactions between bacterial species. This allows us to assess the weight of evidence that inter-species interactions are important in each set of communities. Note that in this neutral model, we expect that real positive correlations are due simply to common species being common together, and that negative interactions are spurious and appear solely due to limitations in sampling (**Fig S8**).

As expected, intestinal communities in the host diverge from the predictions of the neutral sampling model. In all three *C. elegans* strains, we observe more positive correlations than expected (**Fig. 4 G-I**), suggesting that positive interactions among bacteria occurred within the worm host. Consistent with our hypothesis, the significance of this trend is strongest in the *daf-16* strain, where we hypothesized that host control is weakest and interactions among microbes strongest in shaping community composition. The N2 and *daf-16* hosts both show more negative correlations than expected (**Fig. 4D, E**); the trend is significant in *daf-16* communities but not in N2, again consistent with the predictions of our hypothesis. Interestingly, *daf-2* communities show significantly fewer negative correlations than expected (**Fig. 4F**), consistent with the prediction that negative interactions between bacterial species, such as competition, are less important in this apparently stringent host environment.

If the innate immune system actively shapes the microbiome by changing its ecological dynamics, intestinal communities in immune-mutant host strains should show different sensitivities to perturbation as a result. Specifically, we predict that communities in these hosts should react differently to a small change in the relative abundance of species in the metacommunity during community assembly, wherein one bacterial species is “dropped-down” to 10% relative abundance. Here we used a seven-species bacterial metacommunity; MYb56 (*Bacillus*) was removed due to high variation (see also (25)). In these experiments, the “rare” species will experience a drop in expected migration rate into the host (7), resulting in reduced propagule pressure and thus increased variation across hosts in the time to first successful colonization by this species. We chose this setup (as opposed to a larger perturbation such as successive introduction of species) precisely because the mechanism we propose should be sensitive even to a small change in propagule pressure, and because the increased variation in colonization by the “rare” species could increase community variation between individual hosts if priority effects are important (31–33).

We expect this perturbation to have very little effect in a highly stringent host (*daf-2*) where environmental filtering dominates, as we expect bacterial species to colonize the host based on ability to survive this environment rather than on ability to interact with other colonizers. This perturbation may have a large effect in an uncontrolled host (*daf-16*) if there are substantial priority effects (where the early composition of the community affects the species-specific probability of success of later colonists) mediated by inter-species interactions. Alternately, if there are priority effects mediated through changes in the host state rather than through interactions between bacterial species, this perturbation should produce smaller changes in environments where host control is of less relative importance (N2, *daf-16*) than in environments where the host is the dominant factor (*daf-2*).

Our results (**Fig. 5, Table S6**) indicate that host genetics do affect the stability properties of the intestinal community. For N2 and *daf-16* intestinal communities, it is immediately clear that (with the possible exception of the drop-53 condition, **Fig 5B, G**) the perturbation has little to no effect on community composition. These host lineages show a broad but defined range of community compositions when colonized by the full experimental metacommunity (All-7), and the range is essentially recapitulated across conditions. This indicates that these communities are, as suggested previously, converging to a defined range, and that this convergence is stable against the small perturbation applied here.

**Figure 5.**
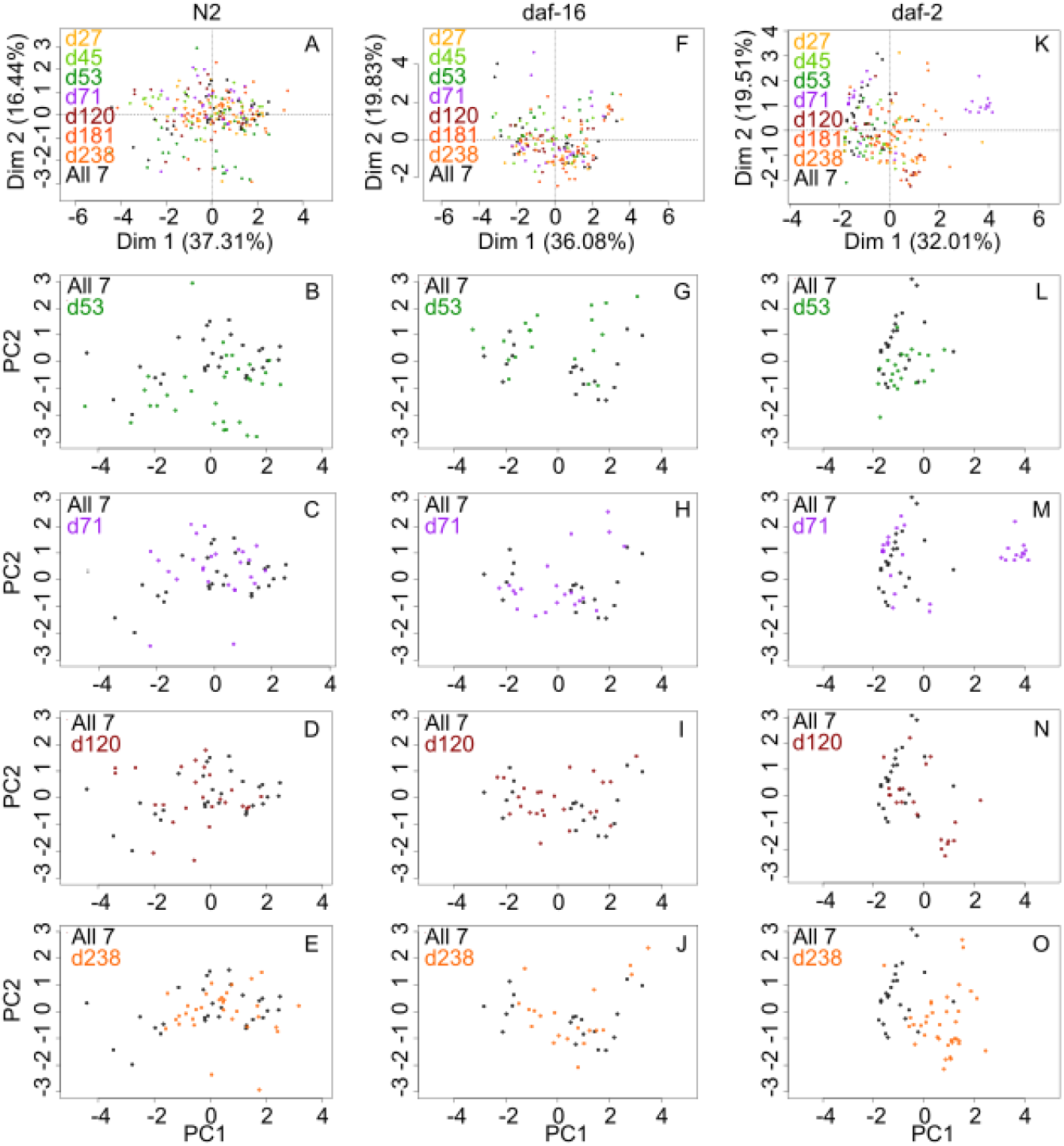
Intestinal bacterial communities in different host lineages react differently to a shared perturbation. In these experiments, adult worms were colonized with an even metacommunity of seven bacterial species (MYb27, 45, 53, 71, 120, 181, 238; All 7) or a metacommunity where each species in turn is dropped (“d”) to 10% relative abundance. Worms were sampled at day 6 of colonization to allow time for communities to pass through early ecological succession. Data represent two (*daf-16*) or three (N2, *daf-2*) independent runs, 12 individual worms per host lineage/metacommunity combination; individual worms with <100 CFU (*daf-2*) or <1000 CFU (N2, *daf-16*) were removed from data to minimize errors due to low colony counts.

By contrast, communities in the *daf-2* intestine (**Fig 5K-O**) occupied a relatively confined ordination space in the All-7 colonization condition, and several of the drop-down colonization conditions (notably drop-71, -120, and -238) resulted in partial or total separation from the All-7 communities. Interestingly, run-to-run variation was considerable in *daf-2* communities but not in N2 (**Fig. S9**). Even within the All-7 condition, it is clear that individual replicates for *daf-2* represent different sub-sets of the total outcome space, while N2-associated communities tend to cover similar ranges across days. In two conditions (drop-71, drop-120), one of three replicates diverged entirely from All-7, and in one condition (drop-238), all three replicates diverged from this baseline. These results are inconsistent with simple sources of experimental error (which would most likely have produced a single divergent run) and indicate that *daf-2* bacterial communities have different stability properties as compared to communities in N2 or *daf-16* hosts. Specifically, *daf-2* communities appear to be both more deterministic (converging to a narrow range of outcomes for a given metacommunity) and more sensitive to initial conditions (such as inevitable run-to-run variation in the metacommunity, **Fig. S10**) than either N2 or *daf-16*. While these experiments are inadequate to fully describe the stability properties of these intestinal communities, these results are consistent with community landscapes for N2 and *daf-16* characterized by wide, stable attractors, while *daf-2* communities appear to occupy a landscape with multiple alternate states (but see Discussion). Further, these results are collectively consistent with the hypothesis that assembly of N2 and *daf-16* communities relies on inter-species interactions among bacteria, while assembly of communities in *daf-2* is controlled largely by the host, and host state changes (plausibly due in part to interactions with bacteria early in colonization) constrain the outcomes of community assembly.

## Discussion

Here we use a tractable model system to demonstrate how host genetics can alter microbiome composition, and how the mechanisms underlying these compositional differences can result in differences in response to perturbation. We observe that changes in *C. elegans* immunity and stress response, in particular in the DAF-2/IGF pathway, are associated with changes in community assembly over time and in stability against perturbations in colonization conditions. These differences in stability result in different possible community states for a given starting condition, indicating that genetic differences in the host can affect the normal operating range and accessibility of alternate states for a microbiome.

These results demonstrate the importance of microbiome ecology for understanding dysbiosis. We observe that host-mediated differences in the ecological drivers of community dynamics result in differences in the availability and accessibility of alternate compositional states. These results suggest that differences in ecological dynamics can produce qualitative differences in propensity toward dysbiosis and the likelihood of return to “normal”. By understanding the drivers of microbiome ecology, it may be possible to gain information about vulnerability to dysbiosis and to predict what types of intervention might be most effective at altering a given community.

It is important to note that these microbiome communities are likely not at a deterministic steady state. First, these are fundamentally not equilibrium systems; these communities are subject to continual disturbance from forces such as migration and physiological shifts in the host (34). Second, even if these communities were heading for a deterministic equilibrium, the relatively short lifespan of the host probably means that intestinal communities are in a transient state (7, 33, 35). Finally, the properties of the host and the intestinal environment are expected to change with age (36–38). Broadly, our results are consistent with previous descriptions of homeorhesis in synthetic communities (39), where a non-stationary system such as a microbiome converges to a trajectory rather than a deterministic point equilibrium, but it is also likely that our communities are in transient states. It is important to consider the underlying dynamics to understand how these results can be generalized, but the conclusions from the data are unaffected - we find distinct and replicable differences in community states and trajectories between different host strains, indicating differences in microbiome ecology driven by host genetics.

Our results are consistent with those of a recent study on host genetics and intestinal community composition in *C. elegans*, but differ in some particulars (25). Both studies indicate that grinder-deficient worms (*tnt-3* in the previous study, *phm-2* here) show increased population sizes in the gut but no significant changes in community composition as compared with N2. Likewise, both studies indicate that all three pathways of innate immunity may be invoked during response of this host to bacteria from its natural habitats. Berg *et al*. observed that communities in *dbl-1* mutants (TGF-β defective) diverged strongly from those in N2, while we saw a smaller effect in this mutant; the previous study focused on differences in *Enterobacteriaceae* abundance, and this clade is not represented in our minimal eight-species community. However, we observed a measurable effect of *dbl-1* even in the absence of *Enterobacteriaceae*, and the previous study did not observe significant community effects in DAF-2/IGF mutants. It is plausible that the different bacterial community and/or the highly controlled liquid culture environment used here (as compared with the experimental soil microcosm approach taken in the previous study) allowed us to detect these genetic effects.

The use of a highly controlled environment carries benefits and drawbacks. We selected a well-controlled, well-mixed environment where worms were grown in liquid medium because it can be replicated accurately, thus allowing us to minimize environmental variation and thereby increase our ability to detect genetic effects. However, as is always the case when a system is abstracted to increase control, this comes at the cost of reducing realism; among other factors, *C. elegans* is not adapted to swimming, and real environments are patchy. It remains unclear how environmental effects might combine with genetic factors to affect the stability properties of host-associated microbial communities. Although there is considerable prior work on the relative contribution of environmental and genetic factors on observed variation in microbiome composition (6, 40–43), at present there is neither theory nor experimental data sufficient for a systems-level explanation of how environment and genetics might act together, or how these interactions can be expected to affect response of microbiomes to perturbation.

There are multiple mechanisms by which DAF-2/IGF signaling might control microbiome composition. This pathway has been implicated in a broad range of stress responses (44, 45) including response to bacterial pathogens, with *daf-2* mutants showing broadly increased resistance (46, 47). Total bacterial colonization can be affected, with *daf-2* mutants showing lower levels of colonization than wild type N2, but increased colonization is not always observed in *daf-16* (48). Some data indicate that DAF-16 is not involved in pathogen response as a primary function but instead maintains a basal level of innate immunity, which regulates response to – and survival in the presence of – non- or minimally pathogenic bacteria such as OP50 (49, 50). However, DAF-2/IGF is an insulin signaling pathway involved in satiety and quiescence behaviors, and we cannot presently rule out the possibility of behavioral differences, such as feeding rate, that might alter bacterial community assembly (51). Further, *E. coli* colonizing DAF-2/IGF mutants show differences in gene expression as compared with those in wild type N2 (52), suggesting that different bacterial factors are required for success; in our multispecies communities, this might result in differences in filtering based on metabolic capacity and/or differences in competitive ability based on metabolic shifts during colonization. Further, we observed that intestinal acidity is an important factor in filtering and control of bacterial competition in *C. elegans* (53), and DAF-16 has been shown to promote acidification of intestinal lysosomes (54); if *daf-2* and *daf-16* mutants are at opposite ends of a spectrum of intestinal acidification, that might explain differences in the stringency of these host environments. Further research, including gene expression analysis in colonized hosts, is necessary to resolve these questions.

## Acknowledgements

This research received no specific grant from any funding agency in the public, commercial, or not-for-profit sectors. This work was supported by funds provided by Emory College.

## Materials and Methods

### Strains and culture conditions

*Arthrobacter aurescens* (MYb27), *Microbacterium oxydans* (MYb45), *Rhodococcus erythropolis PR4* (MYb53), *Bacillus sp. SG20* (MYb56), *Ochrobactrum sp. R-26465 (anthropi*) (MYb71), *Chryseobacterium sp. CHNTR56* (MYb120), *Sphingobacterium faecium* (MYb181), and *Stenotrophomonas* sp. (MYb238) were obtained from the Schulenberg lab (20). Bacterial strains were grown for 48 hours at 25°C with shaking at 300 RPM in individual culture tubes with 1ml of LB. To construct cultures to feed *C. elegans*, appropriate volumes of each strain to attain ~10^8^ CFU/ml were washed in S-medium + 1% AXN, strains were centrifuged 2 minutes at 10K RPM to pellet, and then resuspended in 1ml S-medium + 1% AXN. 100 ml of 100% AXN were prepared by autoclaving 3g yeast extract and 3g soy peptone (Bacto) in 90 ml water, and subsequently adding 1g dextrose, 200μl of 5 mg/ml cholesterol in ethanol, and 10 ml of .5% w/v hemoglobin in 1 mM NaOH.

Laboratory wild-type (N2) *C. elegans* and mutant strains (**Table 1**) were obtained from the *Caenorhabditis* Genetic Center. Nematodes were grown, maintained, and manipulated using standard techniques (55). Briefly, breeding stocks were maintained on NGM plates + OP50 at 25°C (16°C for temperature-sensitive strains AU37 and *glp-4*) and synchronized using a standard bleach/NaOH protocol where eggs were allowed to hatch out in M9 worm buffer overnight (~16h) with shaking (200 RPM) at 25°C. Starved L1 larvae were transferred to 10cm NGM plates containing lawns of *E. coli pos-1* RNAi and incubated at 25°C for 3 days to produce reproductively sterile adults; temperature-sensitive sterile strains AU37 and *glp-4* were grown to adulthood on OP50 under the same conditions. Worms were then transferred to liquid S-medium + 200 μg/ml gentamicin + 50 μg/ml chloramphenicol + 2X heat-killed OP50 (to trigger feeding) for 24 hours, resulting in germ-free adults (**Supp Fig 11**). Adult worms were washed via sucrose floatation before colonization (55).

### Colonization of worms in liquid culture

Bacterial strains were grown separately in 1 mL LB cultures for 48 hours at 25°C and diluted to a constant cell density of 10^8^ CFU/mL. Colonization was performed in well-mixed liquid media according to standard protocols (7) to ensure that all individuals experienced a uniform environment, and had equal access to all potential colonists, for the duration of colonization. Germ-free adult worms were resuspended in S-medium + 1% AXN to a concentration of ~1000 worms/ml. Aliquots of 40μl were pipetted into 96-well deep culture plates (1.2ml well volume, VWR). 20 μl of each bacterial suspension was added to each well (final volume 200 μL). Plates were covered with Breathe Easy sealing membranes and incubated with shaking at 200 RPM at 25°C.

After two days, worms were washed to remove external bacteria and re-fed on a freshly assembled metacommunity; this was done to enforce an evenly distributed community, maximize viability of potential colonists, and minimize bacterial evolution that might lead to unpredictable divergence across replicates. In this step, worms were washed twice in 1ml of M9 Worm Buffer + 0.1% v/v Triton X-100 (M9TX1), rinsed once with S-medium + 1% AXN to remove surfactant, and resuspended in 100 μl of S-medium + 1% AXN. Worms were fed as previously described in a fresh 96-deep well plate.

### Mechanical disruption of colonized worms

Prior to disruption, colonized worms were washed 2X in M9TX1 and moved to an inert food source (heat-killed OP50) for 30 minutes to purge non-adhered bacteria from the gut, then washed and surfaced bleached to remove external bacteria before mechanical disruption of individual worms to retrieve intestinal communities. To clean the exterior cuticle, worms were then rinsed twice with M9TX1, cooled for 15 min at 4°C to stop peristalsis, and bleached for 15 minutes at 4°C with 100μL M9+0.2% v/v commercial bleach. Worms were then rinsed two times with M9TX1 to remove bleach, treated with 100 μl of SDS/DTT solution (965μL M9 + 5 μL SDS + 30 μL 1M DTT) for 20 minutes to permeabilize the worms, and washed once in 1 mL M9. A deep-well plate (2 mL square well plate, Axygen) was prepared by adding ~0.2g of sterile 36-mesh silicon carbide grit (Kramer Industries) and 180 μL of M9TX1 to each well. Worm samples were transferred to a 35mm petri dish with 3 ml of M9TX1, and individual worms were pipetted manually into wells in 20 μL aliquots. The plate was covered with parafilm and kept at 4°C for one hour to reduce heat damage to bacteria. Parafilmed plates were capped with square silicon sealing mats (AxyMat) and disrupted by shaking in a QIAGEN TissueLyser II at 30 hz for 3 minutes. Plates were then centrifuged at 2500xg for 2 minutes to collect all material, resuspended by pipetting, and transferred to 96-well plates for ten-fold serial dilution in PBS (200 μL final volume per well).

### Measurement of bacterial communities

Worm digests were dilution plated onto solid agar for quantification of intestinal communities; bacteria of different types were distinguished on the basis of colony morphology (**Supp Fig 1**). Serial dilutions of 10^−1^ and 10^−2^ of the digests, with the exception of strains DA597 and CB1370, were plated onto modified salt-free nutrient agar (3g yeast extract, 5g peptone, 15g of agar [Bacto] per L). For strain DA597 (*phm-2*), dilutions 10^−2^ and 10^−3^ were plated. Dilutions 10^0^ and 10^−1^ were plated for CB1370 (*daf-2*). In all cases, 100 μL aliquots (of 200 μL total volume) were plated onto individual 10 cm plates using bead shaking. Samples were incubated at 25°C for 48-72 hours to allow distinct morphologies to develop. Colonies were then counted and CFU/worm was calculated for each sample.

### Community analysis, fitting, and ordination

Data analysis was performed in R. Prior to analysis, data were filtered to remove individual worms with <100 CFU/worm (*daf*-2) or <1000 CFU/worm (all other worm strains) to minimize errors from low counts on plates. Bacterial count data were log-transformed and standardized using function *decostand* in *vegan* (56), and principal component analysis was performed using *PCA* from *FactoMineR* (57). Bray-Curtis distances were calculated from log-transformed data using *vegdist* and Shannon diversity was obtained using function *diversity* in *vegan*.

Dirichlet-multinomial fitting and simulations were performed using package *dirmult* (58). Raw data were fitted using the core function *dirmult*, and simulated data were generated using *simpop* with the resulting parameters. To generate simulated data sets for each host lineage, *simpop* was executed with host-specific parameters and a sample size drawn from the vector of CFU/mL totals for that host; each sample in the original data set is simulated this way, resulting in a synthetic data set containing the same number of individual hosts with the same CFU/mL as in the original data. In these simulations, 10,000 data sets were simulated for each host lineage. Spearman correlations were calculated for each real and simulated data set using *rcorr* from the package *Hmisc* (59).

Data for the “drop-down” assays (**Figure 5**) were further filtered before analysis to remove data for bacteria MYb181 from *daf-2* (no counts of this species) and N2 (non-zero counts in 10 of 156 worms), as inclusion of this species resulted in over-weighting of rare MYb181 presence. Communities from *daf-16* worms showed higher prevalence of MYb181 (non-zero counts in 22 of 126 worms), and as removal of this species made no appreciable difference in the qualitative results of the ordination, it was not removed.

